# Variability and uncertainty of data from genotoxicity Test Guidelines: What we know and why it matters

**DOI:** 10.1101/2025.09.24.677002

**Authors:** Giuseppa Raitano, Tessa E. Pronk, Chiara L. Battistelli, Cecilia Bossa, Vasiliki Hatzi, Dimitra Nikolopoulou, Evgenia Chaideftou, Olga Tcheremenskaia, Christelle Adam-Guillermin, Marc Audebert, Birgit Mertens, Martin Paparella

**Affiliations:** Department of Environmental Health Sciences, Istituto di Ricerche Farmacologiche Mario Negri IRCCS, Milano, Italy; KWR Water Research Institute, Nieuwegein, the Netherlands; Environment and Health Department, Istituto Superiore di Sanità, Rome, Italy; Benaki Phytopathological Institute, Laboratory of Toxicological control of Pesticides, Department of Pesticides Control and Phytopharmacy, Athens, Greece; Institute for Medical Biochemistry, Medical University Innsbruck, Austria; Autorité de Sûreté Nucléaire et de Radioprotection (ASNR), PSE-SANTE/SDOS/LMDN, Saint-Paul-Lez-Durance, France; Toxalim, INRAE, INP-ENVT, INP-EI-Purpan, Université de Toulouse Paul Sabatier, Toulouse, France; Scientific Direction of Chemical and Physical Health Risks, Sciensano, Brussels, Belgium

**Keywords:** Genotoxicity, Variability, Uncertainty, OECD, Test Guidelines, Non-Animal-Methods

## Abstract

This review comprehensively examines the variability and uncertainty associated with test guideline (TG)-conform genotoxicity data and explores the respective implications for the integration of non-animal-methods (NAMs) into regulatory frameworks. Historical amendments to OECD TGs are mapped to reveal the method’s evolution that improves the scientific quality of the data but also explains data heterogeneity within available databases. An analysis of the major genotoxicity databases ECVAM, ISSMIC, and OASIS demonstrates substantial variability in genotoxicity calls. Using the EFSA genotoxicity database, which currently harbours the best-curated (meta-) data, we estimate that 22–77% of compounds exhibit similarity of replicate results below 85%, depending on the assay. The potentially most important variables statistically explaining variability and sensitivity were analysed. The practical limitations to identify them with high reliability and to define their optimum needs to be accepted as a qualitative baseline uncertainty. These findings underscore the necessity of contextualizing NAM performance evaluations within the intrinsic variability and uncertainty of animal and in vitro reference data. We propose that this variability is explicitly considered in the development and validation of NAM-based Integrated Approaches for Testing and Assessment (IATAs). This review provides a critical foundation for regulators and scientists aiming to enhance the acceptance and utility of NAMs in genotoxicity assessment.

## 1. Introduction

The thrive towards the reduction and replacement of animal testing for regulatory toxicology has recently gained momentum with the European Commission Initiative for the roadmap towards an animal-free regulatory system (Commission 2024). All current chemical regulations are challenged by this policy, including industrial chemicals, biocides, plant protection products, cosmetics and human as well as veterinary drugs.

In fact, the current dependency of chemical regulation on animal testing appears as an Achilles heel of the European Green Deal and the related Chemical Sustainability and Zero-Pollution Goal. Animal testing is in conflict with all three spheres of sustainability, i.e. economy, society and environment, due to its resource needs in terms of costs and time, due to societal-ethical conflicts and due to uncertainties for species extrapolation and human or environmental variability (Paparella et al. 2024).

In this context, the reduction and replacement of animal tests for genotoxicity assessment of chemicals appears to be a relatively low-hanging fruit. Various in vitro methods are already standardized at OECD level, whilst more are in the pipeline towards OECD standardization (e.g. Toxtracker, in vitro (modified) comet, micronucleus and comet assays in 3D reconstructed skin models, *in vitro* yH2AX/pH3 method). Modern validation relies on the understanding of the mechanisms covered by the model and its human relevance, both of which represent strengths of genotoxicity methods (Hartung et al. 2013). In addition, ample reviews are available on the relevance of the various methods in terms of sensitivity and specificity relative to an agreed reference dataset (Misik et al. 2022).

Some information on the second key validation criterion, i.e. reliability in terms of data variability, is available within OECD validation reports for in vitro and in vivo test guidelines (TGs). Manifold references are provided in the OECD documents (see table S1c), however, as far as available at all, data on variability are not presented following FAIR principles (findable, accessible, interoperable, reusable). Moreover, the limited accessible information appears to be focussed on the variability of the assay readouts (e.g. comet tail length) of negative controls rather than the assay’s results or it is based on very few compounds. No review is available comprehensive for all genotoxicity TGs and assay results for a larger number of compounds. Yet, this information on assay result variability should be key for the regulatory acceptance of experimental and computational non-animal-methods (NAMs) and their use within Integrated Approaches for Testing and Assessment (IATAs). The utility of this information is two-fold. First, it provides key quality criteria for any assay and may be particularly important for an assay aiming to identify substances of high concern with severe regulatory downstream consequences. Second, it shall set expectations for the potential correlation of data from NAMs with data from *in vivo* models. NAMs cannot correlate better with animal reference data than animal reference data may correlate with themselves upon replication. Though this criterion is in theory well respected for computational model validation (no overfitting), a comprehensive retrospective data-based assessment for all TGs was not available to date.

This review aims to provide information on data variability comprehensively, by a) informing on the evolution of OECD TGs over time, as a possible cause for assay results variability within the available databases, b) assessing the constrains of different databases for such retrospective variability assessment in qualitative terms, c) providing a quantitative assessment of TG results variability based on the EFSA genotoxicity database as the one providing most curated data and metadata for applying quality filters, d) exploring potential drivers of variability in quantitative and qualitative terms, and e) contextualizing this uncertainty with current knowledge on uncertainties from other toxicological methods. Finally, we discuss how this review and analysis shall support the development of a NAM-based IATA.

## 2. History of OECD TG changes

### 2.1 Introduction

OECD TGs for genotoxicity testing prioritise essential principles to ensure reliable and relevant outcomes. However, to arrive at this level of scientific quality, over the years, OECD TGs for in vitro and in vivo genotoxicity assays have undergone various revisions and improvements (see Figure 1 and Table S1a and S1b).

**Figure 1.**
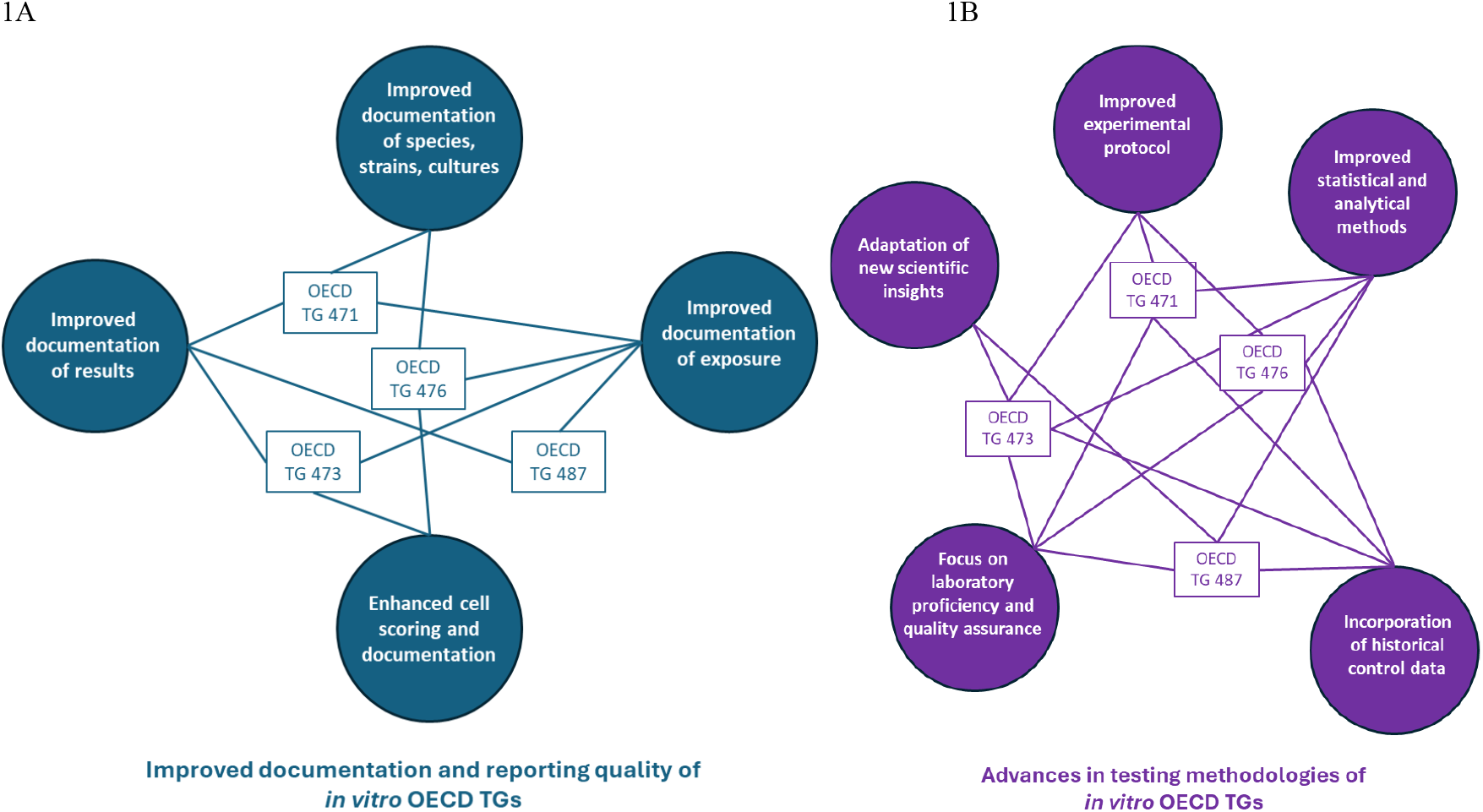
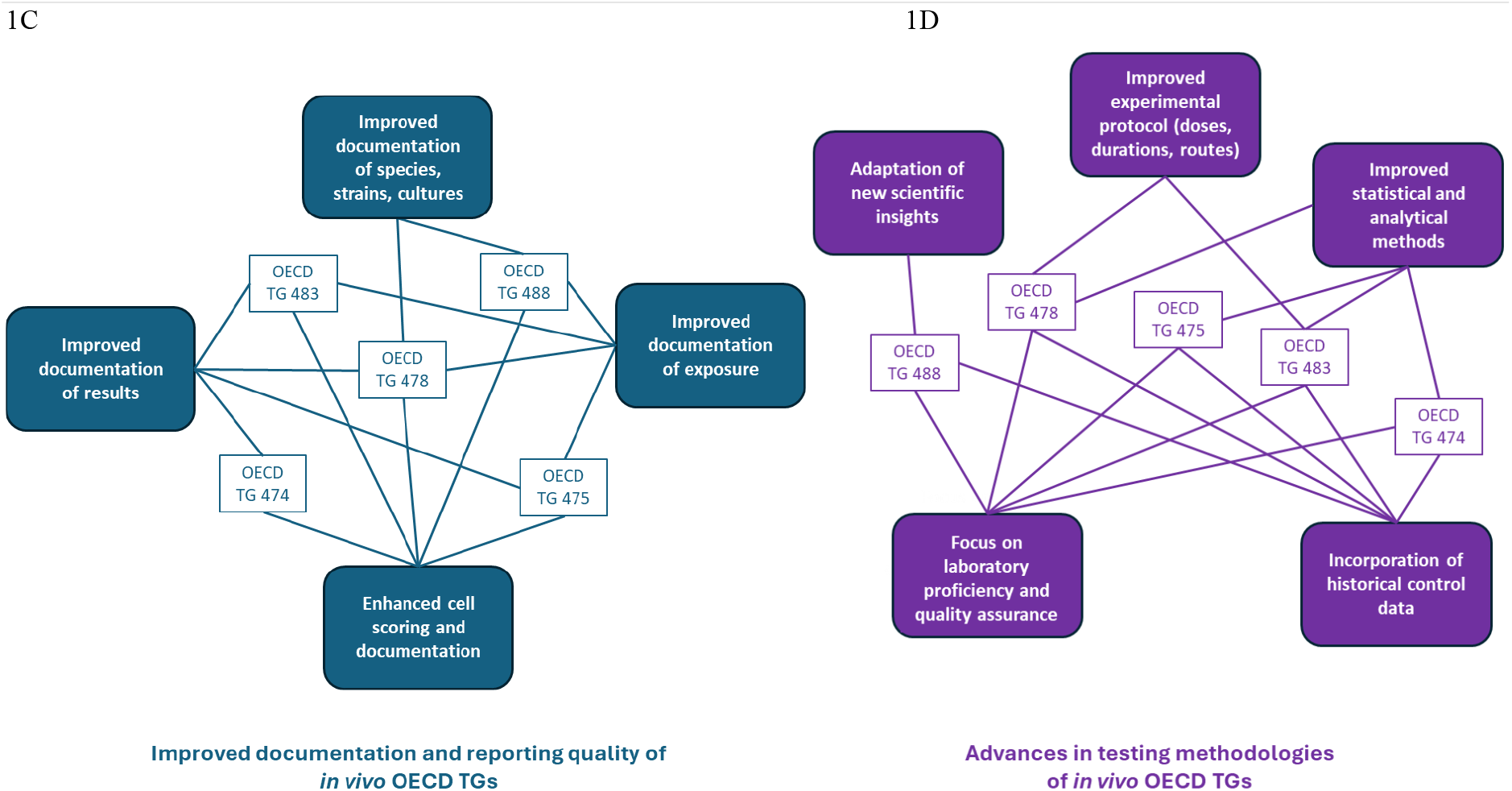
Specifications and amendments of the OECD TGs for in vitro (1A, 1B) and in vivo (1C, 1D) genotoxicity testing from their first implementation till today.

**Figure 2.**
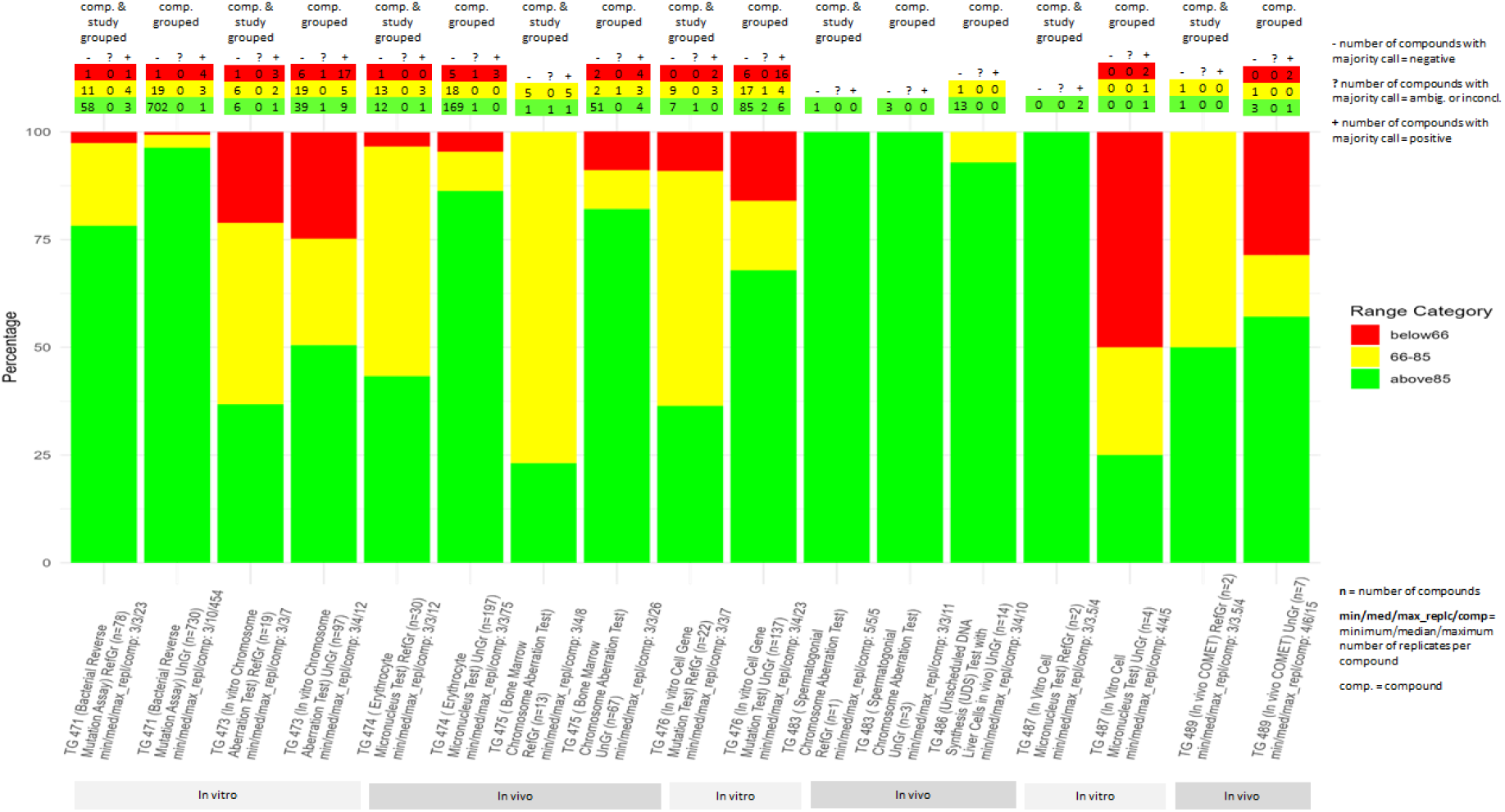
Variability of pseudo-replicates from OECD TG conform genotoxicity studies (see legend and following text for explanation and https://github.com/MartPapa/PARC_genotox_uncertainty for tables including all data)

**Figure 3.**
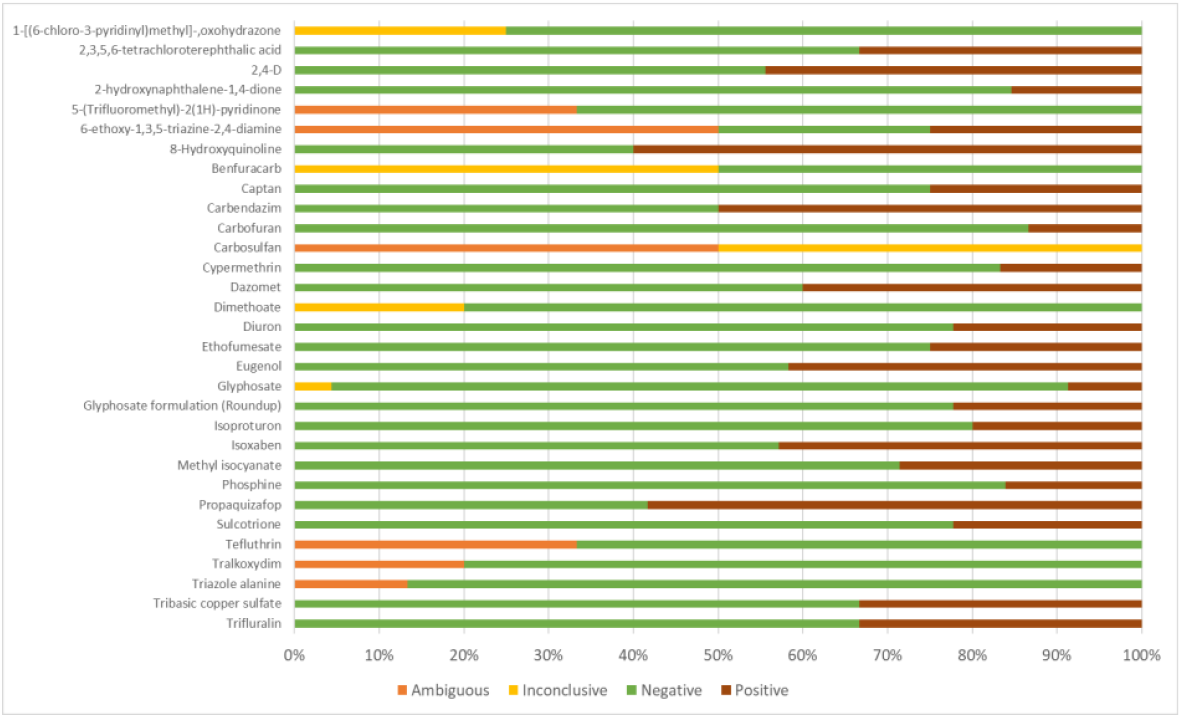
Distribution of the different results for the 31 compounds selected.

In this section, we will explore these changes in the in vitro OECD TGs 471 (Bacterial Reverse Mutation), 473 (Mammalian Chromosomal Aberration), 476 (Mammalian Cell Gene Mutation), 487 (Mammalian Cell Micronucleus), 490 (Mammalian Cell Gene Mutation Test using the Thymidine Kinase Gene) as well in vivo TGs 470 (Mammalian Erythrocyte Pig-a Gene Mutation), 474 (Mammalian Erythrocyte Micronucleus), 475 (Mammalian Bone Marrow Chromosomal Aberration), 478 (Rodent Dominant Lethal); OECD TG 483 (Mammalian Spermatogonial Chromosomal Aberration), 485 (Genetic toxicology, Mouse Heritable Translocation), 488 (Transgenic Rodent Somatic and Germ Cell Gene Mutation), 489 (Mammalian Alkaline Comet Assay).

### 2.2 Method

The methodology followed in this analysis involved a comprehensive review of all OECD TGs relevant to genotoxicity studies. We systematically identified and extracted all revisions and updates made to these TGs over time, drawing from official OECD documentation. Each change was carefully examined to assess its potential impact on test design, data interpretation, and regulatory relevance. To facilitate interpretation, the identified changes were then organized into distinct thematic categories that enabled us to trace the evolution of each TG. By structuring the changes in this way, we aimed to provide a clear and accessible overview of how the OECD TGs have evolved in response to scientific and regulatory advancements. This historical perspective is essential for understanding current data heterogeneity and the implications for comparing legacy and contemporary studies within genotoxicity databases.

### 2.3 Results

An overview of the changes in the OECD TGs of in vitro and in vivo genotoxicity studies is provided below (Figure 1 and Table S1a and S1b).

#### 2.3.1 Improved documentation and reporting standards

Over time revisions across several OECD TGs emphasize *increased documentation and exposure verification* to ensure more reliable and consistent results. For both in vitro and in vivo tests, there is an increasing emphasis on demonstrating *target tissue exposure*, including verification methods. In vitro, these revisions are outlined in *OECD TGs 471, 473, 476, and 487*, while for in vivo tests, they are detailed in *OECD TGs 474, 475, 478, 483, and 488*. Moreover, detailed reporting is now also required for the *species, strain, sex*, and *numbers of animals or cells* used in tests, as well as for statistical measures such as *mean values, standard deviation, confidence intervals*, and other statistical tests applied. Additionally, there is an explicit request for reporting of *cytotoxicity parameters*, and *mutation or aberration types*, ensuring full transparency and replicability of the study.

##### a) Improved documentation of species, strains and cultures

There has been a significant push for *better documentation of species, strains, and cultures* across various OECD TGs. In in vitro testing, compared to the original TG, *OECD TG 471* requires more explicit detailing of the species, strain, and cultures tested (i.e. the inclusion of *E. coli* strains in the Bacterial Reverse Mutation Assay), while *OECD TG 476* mandates more specific information regarding the cell types and cell numbers in the culture/medium per dose. For in vivo testing, the latest version of *OECD TG 488* specifies the need for documenting the *age range of animals* at the start of treatment, as well as a regimen regarding the sex of animals. *OECD TG 478* also calls for more detailed information about the *strain* and *number of animals* used in tests, specifically in accordance with statistical power considerations.

##### b) Improved documentation of exposure

There has been an emphasis on *better documentation of exposure* across both in vitro and in vivo testing. In *in vitro* tests, *OECD TG 471* requires more detailed exposure dose ranges and data on sampling post-application, while both, *OECD TGs 471* and *473*, have updated the exposure route regimen to cover a broader range of cases. *OECD TG 476* also provides more detailed treatment schedules and exposure duration, along with the estimation of the highest exposure dose for different scenarios. Additionally, *OECD TGs 473, 476*, and *487* specify more detailed estimations of the highest exposure dose, including various scenarios. In *in vivo* tests, *OECD TGs 475* and *478* have updated the exposure route regimen to be more detailed and applicable to different scenarios. *OECD TGs 478, 483*, and *488* explicitly require the maximum tolerated dose (MTD) to be included in the dose levels, while *OECD TGs 475, 478*, and *483* call for more detailed exposure dose ranges and sampling post-application data. Furthermore, *OECD TGs 475, 478, 483, 487*, and *488* require verification of exposure to the target tissue, ensuring that exposure is accurately documented across a range of test conditions, which is particularly important to reduce the chance for false negative results.

##### c) Improved cell scoring and documentation

The *cell scoring and documentation* process has also been improved, with more specific requirements for the *treatment, harvesting, sampling*, and *analysis of cells*. The revised OECD TGs emphasize more detailed documentation of cell numbers in both in vitro and in vivo studies. In *in vitro* tests, it is now explicitly required to report not only the number of cells scored, but also the treated and harvested cells as well as the number of cells with chromosomal aberrations (*OECD TG 473*). Additionally, there is a requirement for a comprehensive estimation of cell numbers under various culture conditions or regimens (*OECD TG 476*). For *in vivo* studies, the documentation should include not only the number of cells, but also the number of metaphases scored (*OECD TGs 475 and 483*), and the number of implants, embryos, and other relevant parameters scored (*OECD TG 478*). Updates in in vivo testing to improve the accuracy and reliability of the results, include the use of *automated scoring* (*OECD TG 474*), an *increased number of PCE (polychromatic erythrocytes)* to be screened for *micronucleus (MN)* detection, and an increase in the number of cells needed for the *PCE/NCE (normochromatic erythrocyte) ratio determination*, with a minimum of *20% of control cells* required to assess toxicity accurately (*OECD TG 474*).

##### d) Improved documentation of results

Emphasis has been placed on improving the documentation of results in both in vitro and in vivo OECD TGs. In in vitro tests, the documentation of mutation types (OECD TG 471) and the number and types of aberrations (OECD TG 473) has been requested to ensure more explicit reporting. Additionally, the estimation of the mitotic index (OECD TG 473) and a more detailed estimation of cytotoxicity parameters and indices (OECD TGs 473, 476) have been included. The endpoint has been more clearly defined based on the updated test objective (OECD TG 476), and mutant frequency has now explicitly been requested (OECD TG 476). Furthermore, the documentation of micronuclei/tests (OECD TG 487), and cell lines and cell numbers used in the tests are now more thoroughly documented (OECD TG 487). For in vivo studies, the requirement for more explicit cell and/or centromere scoring and documentation has been incorporated (OECD TGs 483, 475). The number and type of aberrations have to be clearly reported (OECD TGs 483, 475), and the estimation of the mitotic index needs to be provided, where applicable (OECD TGs 483, 475). The analysis for clastogenicity or aneuploidy (OECD TG 474), using techniques such as kinetochore staining with antibodies or DNA probes, has been mandated. Additionally, in vivo documentation now includes more detailed information on species, strain, sex, and animal numbers used in the tests (OECD TG 475), as well as explicit reporting of the maximum tolerated dose (MTD) in the dose levels (OECD TG 475). Finally, the documentation of data related to dose-level frequencies (OECD TG 478) and parameters linked to pre- and post-implantation losses, such as corpora lutea per dam, has been made more detailed (OECD TG 478).

#### 2.3.2 Advances in testing methodologies

Updated protocols have been introduced to expand the exposure dose ranges, durations, and routes of administration in order to accommodate a broader range of test applications. In in vivo testing, an additional requirement has been introduced to document animal body weight and food consumption (OECD TG 478), organ weight data/measurements (OECD TG 483), as well as the explicit reporting of the species, weight variation, and numbers of animals tested (OECD TG 483). These enhancements aim to improve the transparency and reproducibility of the results by offering a more comprehensive description of the test conditions.

##### e) Improved statistical and analytical methods

One notable update is the introduction of trend tests to better detect subtle dose-response relationships, which are now explicitly mentioned in both in vitro (OECD TGs 473, 476, 487) and in vivo (OECD TGs 474, 475, 478, 488) tests. This enhancement improves the sensitivity of statistical analyses, allowing for more accurate detection of potential genotoxic effects at varying doses. Additionally, there is now an increased requirement for the documentation of mean values and standard deviations in the reported parameters, with specific guidelines for in vitro tests (OECD TGs 476, 487) and in vivo tests (OECD TGs 475, 478, 483). These updates also include the need for detailed dose-response data, including maximum tolerated dose (MTD) (OECD TGs 475, 478, 483, 488) considerations, further refining the accuracy and relevance of the data.

##### f) Incorporation of historical control data

The use of both *negative and positive historical control data* allows researchers to better assess the reliability of their findings. This approach is emphasized across both in vitro and in vivo guidelines to strengthen data accuracy. Additionally, there is a strong push for *transparency* and *reproducibility*, with detailed data records and methodologies that ensure studies can be independently verified. Relevant OECD TGs highlighting the need for this data include *471, 473, 476, and 487* (in vitro) and *474, 475, 478, 483, and 488* (in vivo).

##### g) Focus on laboratory proficiency and quality assurance

A key area of focus has been the increased *requirements for laboratory proficiency*, ensuring that laboratories demonstrate consistent performance in generating reliable data. This includes greater emphasis on *protocol standardization* and adherence to detailed *OECD TGs* to ensure that testing is conducted in a reproducible and standardized manner across laboratories. The importance of documenting laboratory proficiency has been highlighted, with *OECD TGs 471, 473, 476, and 487* (in vitro) and *OECD TGs 474, 475, 478, 483*, and 488 (in vivo) specifically requiring verifiable proficiency in conducting tests.

##### h) Adaptations for new scientific insights

With the rapid advancements in scientific knowledge, *methodologies for clastogenicity, aneuploidy*, and *cytotoxicity* assessments have been significantly improved (e.g. *OECD TGs 473, 487*). In addition, *detailed guidelines* have been established for *sampling post-treatment* and *analysing key reproductive or genetic endpoints*, helping researchers link their data to potential health outcomes, including reproductive health and genetic mutations. The sampling of *reproductive tracts* for *sperm collection* and revised guidelines on the *timing* for *rodent spermatogonial stem cells* to mature into sperm and reach the *cauda epididymis have been documented* (in *OECD TG 488*).

##### i) Expanded applicability, reliability and relevance

The *objectives and endpoints* of testing have been refined to align with *updated scientific knowledge*, allowing for more reliable and relevant findings. There is also a stronger emphasis on the *sensitivity* and *specificity* of tests, particularly in detecting various types of genetic damage. Furthermore, *advanced staining* and *detection techniques*, such as *kinetochore staining* for detecting *aneuploidy* in the *in vitro mammalian cell micronucleus assay (OECD TG 487*), have been introduced to further enhance the accuracy of the results.

### 2.4 Conclusion

Over the years, the improvements to the OECD TGs for in vitro and in vivo genotoxicity assays reflect a commitment to modernizing the testing framework through comprehensive updates to testing methodologies, data documentation, and statistical analysis. These enhancements ensure the precision, transparency, and thoroughness required for reliable safety assessments, aligning with the latest scientific developments and significantly improving the relevance and reliability of genotoxicity studies in both regulatory and research contexts. Specifically, the improvements include testing at a minimum of three doses or concentrations, ensuring exposure within the toxic range, and utilizing both positive and negative controls. It is necessary to provide evidence of the test substance’s presence at the target cells or tissues, with appropriate timeframes for sample collection. Key aspects include documented statistics such as mean values, standard deviations, and confidence intervals, alongside biological relevance with respect to exposure routes, treatment durations, and the consideration of both specific genotoxic effects and potential non-specific cytotoxicity. Important updates to the guidelines involve detailed documentation of the number of animals used, their species and strain, sex, age range, routes of administration, and critical parameters such as food consumption and body or organ weights. There is also a greater emphasis on specifying types of mutations and chromosome aberrations, as well as cytotoxicity parameters and the maximum tolerated dose (MTD). The inclusion of historical control data, enhancements in staining and detection methodologies, and the proficiency of the testing laboratories further enhance the overall scientific quality of studies.

However, the continuous adaptations of OECD TGs over time are reflected as a heterogeneity of data-quality within the currently available databases. Different databases allow data selection based on a different level of detail on their specific quality (see section 3). Yet, for a statistically meaningful analysis of data-variability it is practically unavoidable to include also data from earlier than the latest version of the TG. This represents an uncertainty in the analysis of the methods data variability.

## 3. Available knowledge of variability between databases

### 3.1 Introduction

Different databases may provide method-specific genotoxicity calls that differ for the same compound. This represents an uncertainty for the assessment of the variability of genotoxicity data from similar methods. To characterize this uncertainty, an analysis of these differences between databases is provided. Existing public databases containing in vivo micronucleus test results were compared; specifically, the “EURL ECVAM Genotoxicity & Carcinogenicity”(Corvi and Madia 2018; Madia et al. 2020), the “in vivo mutagenicity (micronucleus test) ISSMIC” database (Benigni et al. 2021) and the Micronucleus OASIS database (OECD 2025). Starting from these databases, three datasets were built, which include chemical substances with in vivo micronucleus results and chemical identifiers (chemical name, CAS number and/or SMILES code). An analysis is provided for the three different data sets in terms of compound coverage, differences in concluding on genotoxicity calls, and the concordance of these micronucleus calls for identical compounds between different datasets.

### 3.2 Results

The Genotoxicity & Carcinogenicity ECVAM database (hereafter called ECVAM) was developed by the European Reference Laboratory for Alternatives to Animal Testing (EURL ECVAM) and contains data of multiple genotoxicity assays from different sources together with carcinogenicity results. ECVAM was built from the database collecting data of Ames positive substances (Kirkland et al. 2014a; Kirkland et al. 2014b), later expanded to include data on Ames negative substances as well (Madia et al. 2020). The ECVAM dataset used for the analysis was built including ECVAM substances with in vivo micronucleus assay results, taken from the list of both Ames positive and negative chemicals. In the original ECVAM database, the results for micronucleus data are categorized as follows: Positive, Weak positive, Negative, Equivocal and Inconclusive results; the criteria for the assignment of the overall call, described in Kirkland et al 2014b and Madia et al 2020, are summarized in table 2. The original list has been cleaned by eliminating substances without a chemical identifier (CAS number or SMILES) and resolving duplicated IDs. The results reported in the original database were grouped into 4 categories, depending on the outcome, as follows: Positive (including positive, weak positive and in the presence of at least one positive result), Negative, Equivocal and Inconclusive results (the latter reflecting the original categories). The resulting dataset consists of 608 unique chemicals with in vivo micronucleus data. Details of ECVAM for in vivo Micronucleus data used in this analysis are reported in table 1.

**Table 1:** main characteristics of the datasets for in vivo micronucleus data employed in this analysis. Pos: Positive; Neg: Negative; Eq: Equivocal; Inc: Inconclusive; PB/BM: peripheral blood/bone marrow.

| **In vivo Micronucleus datasets** | **Total number of chemicals** | **Overall call (N chem.)** | **Experimental details** |
| --- | --- | --- | --- |
| **ECVAM** | 608 | Pos (247); Neg (280); Eq (73); Inc (8) | rat/mouse, PB/BM |
| **ISSMIC** | 562 | Pos (190); Neg (97); Eq (48); Inc (227) | rat/mouse, Male/female<br>Admin. Route, PB/BM, Toxicity at the target |
| **OASIS** | 557 | Pos (258); Neg (299) | human/rat/mouse/mammals/rabbit<br>PB/BM |

**Table 2:** criteria for the overall call definition within in vivo micronucleus databases of this analysis.

| **Databases** | **Positive Overall** | **Negative Overall** | **Inconclusive Overall** | **Equivocal Overall** |
| --- | --- | --- | --- | --- |
| **ECVAM** | i) Clear evidence of a positive response from a single or more studies; or<br>ii) Positive response in one species or sex and negative in the other but clear evidence that systemic exposures were greater in the positive than in the negative study | i) No evidence of a positive or equivocal response and OECD TG or recommended best practices fulfilled; and<br>(ii) Some direct or indirect evidence that the test substance reached the target tissue, otherwise it was considered inconclusive | i) No definitive conclusion in term of compliance with the requirements of the current OECD TGs or recommended best practices; or<br>ii) No evidence that adequate levels of toxicity were achieved (in vitro); or<br>iii) No proof of target cell exposure (in vivo); or<br>iii) Result not obtained via adequate testing, i.e., "no valid data". | (i) Ambiguous, doubtful, questionable, or inconsistent (e.g., a positive and a negative test) results within a study; or<br>(ii) A dose-related increase in effects, close to the borderline of biological significance, but the responses not biologically and/or statistically significant, and no independent repeat experiment available to verify the response; or<br>iii) Some evidence of a positive response that could not be dismissed, but no consistent responses in the same test system across different studies; or<br>iv) both positive and negative findings across different studies of apparently equal validity, and no clear positive or negative overall call possible based on the weight of evidence |
| **ISSMIC** | Positive response in at least one experimental group (mouse, rat, male and female) | Negative response (no induction of micronuclei) with clear evidence of cell exposure | (i) No induction of micronucleus formation nor target cells toxicity and<br>(ii) No induction of micronucleus formation and no target cell toxicity information available in the literature | Equivocal in at least one experimental group, and negative in the other experimental groups |
| **OASIS** | Criteria for the overall calculation, in case of multiple experiments on the same endpoint, were not reported |  |  |  |

The ISSMIC database, was developed by Istituto Superiore di Sanità (Benigni et al. 2012) and contains results relative to the in vivo micronucleus mutagenicity assay for 566 chemicals. Standard chemical data fields are included together with biological data fields: outcomes in bone marrow cells, peripheral blood cells and splenocytes in four experimental groups (mouse, rat, male and female); details such as toxicity biomarker (target cell toxicity). During the development of the database, the authors paid particular attention to the definition of negative results: the target cell toxicity was used as demonstration of the chemical’s interaction with target cells, and for each experiment, this information was included in the database. ISSMIC results are stratified into the categories of (i) Positives (n = 190), (ii) Equivocal (n = 48), (iii) Negatives with clear evidence of cell exposure (n = 97) and (iv) Inconclusives (n = 231). Inconclusive results include (i) chemicals that neither induce micronuclei nor target cell toxicity and (ii) chemicals that do not induce micronuclei and for which target cell toxicity information is not available in the literature. While the ISSMIC database is available as a source of data in the QSAR Toolbox, the stand-alone version (https://www.iss.it/en/isstox) was used for the present analysis. The general characteristics of ISSMIC dataset used in this analysis are reported in table 1.

The Micronucleus OASIS database (hereafter called OASIS) was developed and maintained by Laboratory of Mathematical Chemistry (LMC) in Burgas (Bulgaria). OASIS public version, available in the QSAR Toolbox, contains results relative to the in vivo micronucleus assay for 557 chemicals, containing predominantly in vivo bone marrow and peripheral blood micronucleus data for rats and mice. Results included in OASIS were collected from various sources, including articles and public websites, whose reference is reported in the metadata of each individual chemical. The outcomes are categorized in (i) Positive or (ii) Negative results. In the present analysis, the QSAR Toolbox 4.6. (OECD 2025) implementation for OASIS database was used. The general characteristics of the OASIS dataset are reported in table 1.

The overlap between the different database pairs was analysed, and for each pair, the concordance on the chemicals in common was calculated. The concordance rate (as reported in table 3) has been calculated by adding the counts of the concordant results in each pair divided by the total number of substances in common.

**Table 3:** concordance (%) calculated for each pair of databases, on the common substance (the number of substances is shown in brackets) with available experimental results on in vivo micronucleus test.

| **DB comparisons** | **Concordance** |
| --- | --- |
| **ECVAM vs. ISSMIC** | 50% (of 227 substances) |
| **ECVAM vs. OASIS** | 75% (of 286 substances) |
| **OASIS vs. ISSMIC** | 43% (of 207 substances) |

Regarding the ISSMIC/OASIS pair, the overlap between the datasets involves 207 substances. Table 4 summarizes the comparison of the overall calls for the overlapping chemicals. Marked differences between the two datasets appear both for the categories of substances not present in OASIS (i.e., Equivocal and Inconclusive results) and in the classification of Positive and Negative results. This is reflected in a concordance between the two datasets of only 43% (table 3). This could partly be due to different micronucleus data (going into the details of this possibility would require re-examining all the substances, one by one, and is beyond the scope of this analysis) or the way of interpreting the data in terms of an overall call.

**Table 4:**
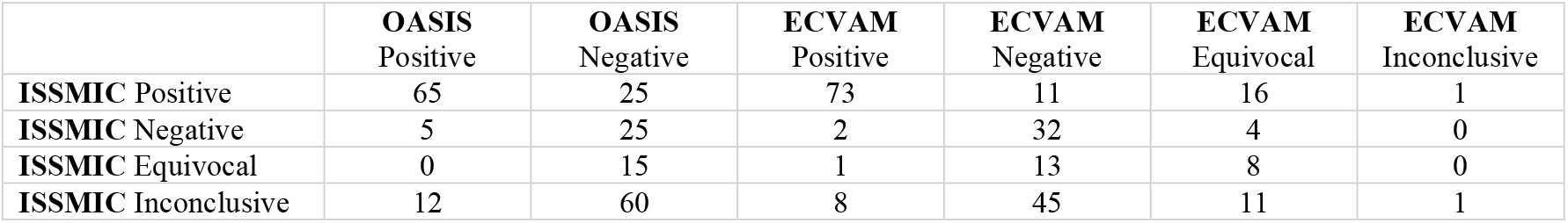
distribution among the different overall calls of ISSMIC, OASIS and ECVAM for the 207 and 227 overlapping chemicals, respectively.

ISSMIC and ECVAM share 227 substances. Even in this case, the concordance of the results is quite poor (i.e., 50%, table 3). This is, at least partly, clearly due to differences in the way of concluding on the overall call (table 2). Indeed, the ISSMIC criteria are decided a priori (e.g., the overall is assigned positive if it is positive in at least one experimental group) while in the ECVAM database, the authors assigned the overall call by weighing the evidence, on a “case by case” basis. Moreover, similarly to the previous case with OASIS, also in this comparison, it is observed that the substances classified as inconclusive for ISSMIC are negative for ECVAM. This highlights a significant discrepancy in taking into account parameters related to target toxicity. Finally, the ECVAM/OASIS pair includes 286 common chemicals: table 5 summarizes the distribution among the different overall of these overlapping chemicals. In this case, the concordance displays a higher value (i.e., 75%, table 3). Still, a non-negligible percentage of chemicals shows a different outcome depending on the provenance database.

**Table 5:** distribution among the different overall calls of OASIS and ECVAM datasets, on the 286 overlapping chemicals.

|  | **OASIS Positive** | **OASIS Negative** |
| --- | --- | --- |
| **ECVAM Positive** | 126 | 17 |
| **ECVAM Negative** | 8 | 89 |
| **ECVAM Equivocal** | 26 | 19 |
| **ECVAM Inconclusive** | 1 | 0 |

**Table 6.** Compounds for which more than one variable is relevant. InvivoTexp = INVIVOTISSUEEXP; N_ind = NUMBER_INDIVIDUALS; LR = literature_reference.

| Name of compound | variable n. 1 | variable n. 2 | variable n. 3 | variable n. 4 | variable n. 5 |
| --- | --- | --- | --- | --- | --- |
| 1-[(6-chloro-3-pyridinyl)methyl]-,oxohydrazone | strain | sex |  |  |  |
| 8-Hydroxyquinoline | strain | N_ind | InvivoTexp | LR |  |
| Benfuracarb | InvivoTexp | LR |  |  |  |
| Captan | N_ind | sex | InvivoTexp |  |  |
| Carbofuran | strain | LR |  |  |  |
| Carbosulfan | strain | N_ind | sex | InvivoTexp | LR |
| Dazomet | N_ind | sex | route | InvivoTexp |  |
| Dimethoate | strain | N_ind | sex | route | InvivoTexp |
| Diuron | strain | N_ind |  |  |  |
| Phosphine | sex | InvivoTexp |  |  |  |
| Propaquizafop | route | InvivoTexp |  |  |  |
| Sulcotrione | strain | sex |  |  |  |
| Triazole alanine | species | sex |  |  |  |

### 3.3 Conclusion

Different databases include an overlapping set of compounds. For a large proportion thereof, i.e. 25% to 57% for the compounds with in vivo micronucleus data analysed here, the method-specific genotoxicity calls may differ. This is partly due to the database-specific approaches for concluding on these calls, which may be strict a priori rules or expert-based evaluations. Furthermore, the variability is probably also due to the presence of data from different studies of differing quality and protocols (see section 2). The latter cannot be easily analysed, since the databases investigated here also differ in the availability of metadata reported. This limitation is important to be recognized by regulators that may use different genotoxicity databases, including the QSAR toolbox, for the assessment of compounds as such or within grouping and read-across approaches.

Moreover, for deciding on the regulatory acceptability of a QSAR model, and on the reliability of the resulting predictions, its predictive power must be compared with the intra- and inter-laboratory reproducibility of experimental replicates. Yet, for a systematic analysis of possible experimental variability of genotoxicity data from similar methods, a database including rich, curated metadata is essential. Recently, the latter has become available as the EFSA genotoxicity database, which is already included in the OECD QSAR toolbox.

## 4. Improving Genotoxicity databases

Besides the databases already described under Sections 3, there are ongoing efforts to enhance the data range, both in terms of chemical space and covering test types especially for those with a limited amount of data. For instance, EFSA has launched two projects with the purpose of extension and update of the existing EFSA genotoxicity database (Metruccio et al. 2017a; Metruccio et al. 2017b).

The original EFSA genotoxicity database was commissioned by EFSA in 2014 after the EFSA panel on Plant Protection Products and their Residues issued a scientific opinion (EFSA-PPR 2012), which included (Q)SAR modelling as part of an integrated approach to assess genotoxicity of pesticide active substances and their metabolites. The database was finalised in 2017 and includes the genotoxicity data for about 380 active substances and 600 metabolites. Subsequently, this database was used to evaluate the model performance of (Q)SAR models for five in vitro and in vivo genotoxicity endpoints (Benigni et al. 2019): Bacterial Reverse Mutation Assay (Ames test), Mammalian Bone Marrow Chromosome Aberration Test, Mammalian Erythrocyte Micronucleus Test, in vitro Mammalian Chromosome Aberration Test, and in vitro Mammalian Cell Gene Mutation Test. Statistically reliable results were obtained only for the Ames test, while the reliability of prediction for other genotoxicity endpoints was not sufficient, underlying the need for further data collection.

A recently published external scientific report of EFSA “Extension of the EFSA Pesticides Genotoxicity Database” by German Federal Institute for Risk Assessment (BfR) (BfR 2025; Foil et al. 2025) describes the recent project, in which the Ames and in vitro micronucleus datasets were transferred from the original MySQL BfR Genotoxicity Database to IUCLID. The original studies’ reports were re-evaluated according to the criteria set in OECD test guidelines (TG) 487 In Vitro Mammalian Cell Micronucleus Test and 471 Bacterial Reverse Mutation Test. This extension of EFSA genotoxicity database includes 82,082 single data, 9,303 incubations, 481 reports, and 349 substances for the Ames test and 3,844 single data, 685 incubations, 203 reports, and 183 substances for the in vitro micronucleus test.

The ongoing project “Update of the EFSA Pesticides Genotoxicity Database” (Novello et al. 2024) is a 3-year collaborative effort developed by Istituto Superiore di Sanità (ISS) and Innovatune SRL (https://www.innovatune.com/). The updated database will be published as external scientific report of EFSA in the next months and will include the update of the currently available EFSA genotoxicity database for about 140 new pesticides active substances and about 280 metabolites both in Excel and IUCLID formats.

OpeFoodTox is another EFSA database that includes the genotoxicity data. The database is currently undergoing an update that will be completed in 2026 (Carnesecchi et al. 2023; Iovine et al. 2025). It is a structured database summarising the outcomes of hazard identification and characterisation for the human health (all regulated products and contaminants), the animal health (feed additives, pesticides and contaminants) and the environment (feed additives and pesticides).

In summary, the need for enhanced genotoxicity database development, including curated data and meta-data, from established as well as new methods, for a broader set of compounds, is well recognized and respective work is in progress.

## 5. Available quantitative knowledge about the variability of pseudo-replicate data from OECD TG conform studies

### 5.1 Introduction

The latest publicly available EFSA Genotoxicity database is used for an analysis of data variability. This database provides the most detailed meta-data for applying quality filters. The intention of this analysis is to provide an estimate for regulators, how variable the data from TG conform studies may be. It contains the caveat that the database builds on EFSA opinions between 2004 and 2016 and these include older data, which were acceptable for regulation at the date of the opinion. Moreover, it is noted that the TGs are not Standard Operating Procedures (SOPs) and allow some variants of the test protocol and all TG conform studies are generally considered acceptable. Here we use the term “pseudo-replicates” for replicates which may have been generated with different protocol variants acceptable though within the frame of the OECD TG (e.g. species, exposure route, solvent used, exposure and sampling regime, target cells analysed), at the time of the EFSA opinions.

Regulators usually receive no or very few pseudo-replicate studies, such that the true data variability in TG conform studies remains practically hidden to them. Therefore, the variability estimate provided in this section represents data uncertainty for regulators.

### 5.2 Method

For this analysis, a special version of the EFSA genotoxicity database (EFSA 2017) was used, which included study-codes for each line, such that results can be grouped for studies. The literature study underlying the study code remained confidential.

The following filter criteria were applied to the database:

1. guideline_qualifier = “According to” or “Equivalent or similar to” the TG version acceptable at time point of the regulatory submission and EFSA opinion
2. acceptability = “acceptable”
3. genotoxguidelines = OECD *TGs 471*|*473*|*476*|*487*|*490*|*474*|*475*|*483*|*486*|*488*|*489*
4. entries with at least 3 (pseudo-) replicates (either within studies or between studies, see below for A and B)
5. use of compound name for analysis (no filtering for substance name and/or qualifier): The compound name relates to the item tested, whereas the substance name relates to the substance assessed. The compound may be identical to the substance, or a metabolite of the substance or part of a mixture in a group or mixture assessment. For the data-variability analysis, we were just interested in the variability of the data from the compounds tested. It is noted that a complete and correct removal of duplicates is not possible with this publicly available database, since not all test-variables are presented in the database (as columns). Therefore, the impact of removing or not the duplicates as far as possible with this database was analysed. The impact was minimal and visible only for the *TG 489* (in vivo comet assay), where 7 versus 4 compounds with at least 3 replicates remained after filtering. Moreover, we were informed by experts with access to the full confidential database, that the removal of these three specific entries would not be correct due to some test-variables not visible in the database. Therefore, we finally did not remove any seemingly duplicated lines.

Thereafter, two approaches for analysis were taken:

A. The data were grouped for compound names (com_name). This analysis results in an estimate for data-variability *within* studies, that may result e.g. from measuring within one study different post-exposure sampling times, sex, different microbial strains, with or without S9 mix and else.
B. The data were grouped for compound names and studies (com_name, literature_reference). This analysis results in an estimate for data-variability *between* studies. The following conclusion is taken at the study level: If all results within the study are negative the result is considered negative, if one result is positive, the result is considered positive, otherwise (i.e. only negative and/or inconclusive/ambiguous results are available) the result is considered inconclusive/ambiguous.

With both approaches, A and B, the data were extracted to derive the:

- number of pseudo-replicates per compound
- minimum, median, maximum number of pseudo-replicates per compound per TG
- % of chemicals with a <66% probability for identical results (either positive, negative or inconclusive/ambiguous) upon replication,
- % chemicals with a 66-85% probability for identical results
- % chemicals with >85% probability for identical results
- majority call for each compound, i.e. if majority of pseudo-replicate results indicated “negative”, “ambiguous or inconclusive” or “positive”
- number of compounds with a majority call for negative, ambiguous/inconclusive or positive result for each of the three categories (< 66%, 66-85%, > 85% of identical results from pseudo-replicates)

The boundaries of the three categories (< 66%, 66-85%, > 85% of identical results from pseudo-replicates) were selected based on the following arguments:

✓ a 66% probability for an identical result means that it is twice as likely to receive the same result as not
✓ 86% probability for an identical result means that it is about 6 times more likely to receive the same results than not

The R-code for this analysis, including the EFSA database file with the literature codes is available via Github (https://github.com/MartPapa/PARC_genotox_uncertainty).

To what extent data variability may be explained by variables within TG conform test protocol is explained in the next section 6.

### 5.3 Result and Conclusion

Grouping results just by compound rather than by compound and study indicates a lower number of compounds within the very low (<66%, red in figure) and low (66-85%, yellow in figure) similarity range. This appears to indicate that few positive or ambiguous/inconclusive test-results drive the variability much more for regulatory study results, where a single positive testresult within the study triggers a conclusion for an overall positive hit call for the compound. However, since variability from this regulatory data interpretation is most relevant for the purpose of this article, in the following, we focus on the latter, i.e. the variability of study results per compound.

The number of compounds with at least three pseudo-replicate data ranged from 78 for *TG 471* (Ames), 30 for TG 474 (in vivo micronucleus), 22 for *TG 476* (in vitro gene mutation), 19 for *TG 473* (in vitro chromosomal aberration), 13 for *TG 475* (in vivo bone marrow chromosomal aberration), 2 for *TG 487* (in vitro micronucleus) as well as for *TG 489* (in vivo comet) and 1 for *TG 483* (in vivo spermatogonia). The number of replicates per compound was minimal 3, on average (median) 3 to 5 and maximal 23 for each of the TGs, which are analysed here.

The percentage of compounds with pseudo-replicate similarity at the study level in the very low (red in figure) or low (yellow in figure) range, is distributed between 22% for *TG 471* (Ames), 56% for *TG 474* (in vivo micronucleus), 63% for *TG 473* (in vitro chromosomal aberration), 64% for *TG 476* (in vitro gene mutation) and 77% for *TG 475* (in vivo bone marrow chromosomal aberration). For all these TGs, the proportion of compounds with a positive or ambiguous/inconclusive majority call (at study level) is higher within the very low (red) or low (yellow) similarity range.

For the other TGs, fewer than 3 compounds are available with at least 3 pseudo-replicates, such that an interpretation of these data is highly uncertain: *TG 487* (in vitro micronucleus test), *TG 489* (in vivo comet), and *TG 483* (in vivo spermatogonia test).

## 6. OECD TG conform study-variables that may explain the data-variability

A multivariate analysis was carried out with the aim of identifying possible drivers of data variability, in order to potentially develop recommendations to reduce the variability of data from genotoxicity methods.

This exploratory analysis, closely linked to the specific composition of the dataset used, made it possible to identify the most relevant variables for the studied phenomenon (multiple results), facilitating an understanding of the data structure and selection of the most meaningful features.

We used the Mammalian Erythrocyte Micronucleus Test (*TG 474*) as a paradigm for the analysis, as it was the most populated dataset (944 entries for 197 individual compounds), according to the selection criteria explained above (approach A, in section 5.2., filtered input-data see supplement 2a), with the highest percentage of compounds in the very low (red in figure) or low (yellow in figure) range. In the Supplement 2b, we reported the same analysis performed for the other three most populated datasets among the in vivo and in vitro methods (i.e. Mammalian Bone Marrow Chromosome Aberration Test (*TG 475*), in vitro Mammalian Chromosome Aberration Test (*TG 473*) and in vitro Mammalian Cell Gene Mutation Test (*TG 476*)).

Of the 197 individual compounds, 31 had Mammalian Erythrocyte Micronucleus Test data with multiple results (equivocal, inconclusive, negative and positive outcomes).

### 6.1 Method

Genotoxicity studies often involve analysing large data sets with many variables (e.g. different chemicals, gene expression levels, etc.). Random forest (RF) technique (figure 4), among the other multivariate methods, can efficiently handle such high-dimensional data without overfitting, a common problem with simpler models. Furthermore, RF offers numerous advantages in the exploratory phases of data analysis. It makes it easy to identify which characteristics (features) are most relevant for explaining variability in the data. This improves understanding of the dataset’s structure and helps select the most significant variables for subsequent analysis or building simpler models. The algorithm is robust in the presence of noisy data and outliers and can effectively handle missing data while maintaining good performance and reliability in exploratory analyses.

**Figure 4.**
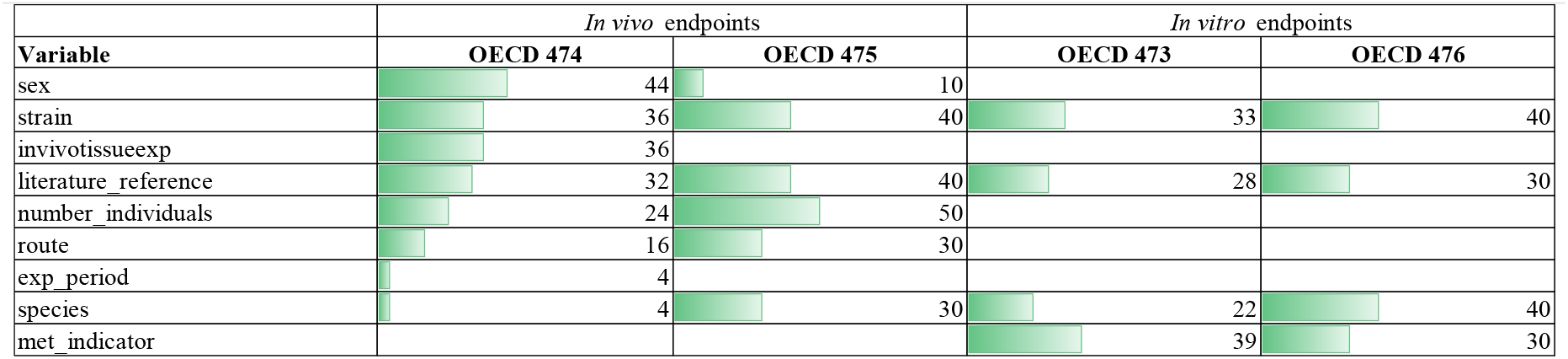
Distribution of the variables and their percentage coverage across the four datasets.

**Figure 5.**
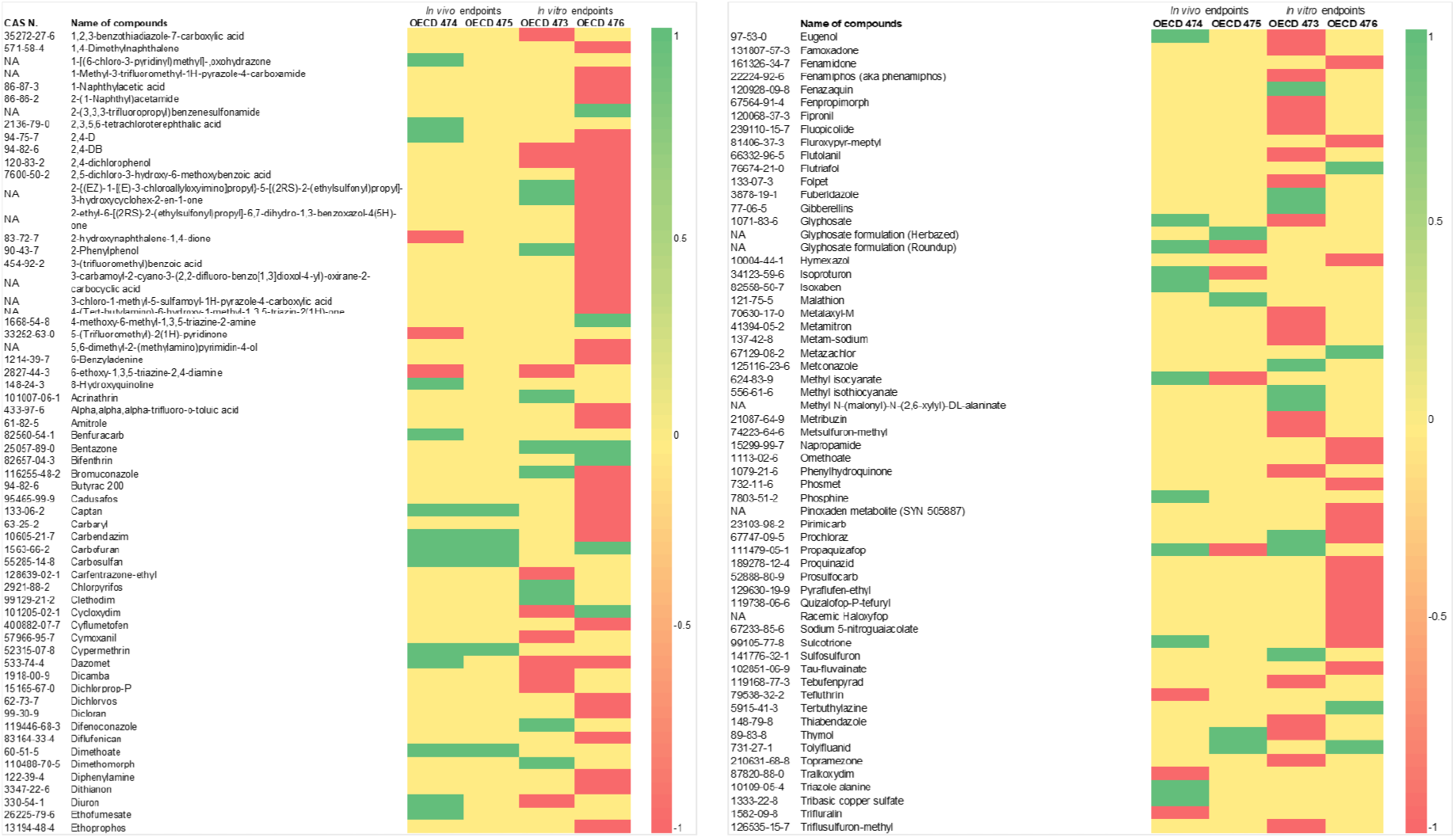
Heatmap representing the 122 compounds belonging to all four datasets. RED=with multiple results but not explained by the RF; GREEN= with multiple results, explained by the RF; Yellow= non present

The RF technique applied consists of a selected number of decision trees. Each of the decision tree models is trained on a different set of rows (records) and a different set of columns (describing attributes), where the latter can also be a bit-vector or byte-vector descriptor (e.g. molecular fingerprint). The row sets for each decision tree are generated by bootstrapping and have the same size as the original input table. For each node of a decision tree, a new set of attributes is determined by taking a random sample of size sqrt(m), where m is the total number of attributes. The output model describes a random forest and is applied in the corresponding predictor node using simple majority voting. Due to the combination of multiple trees and the random selection of data and variables, RF reduces the risk of overfitting compared to single decision trees, providing more generalisable results, even in the exploratory phase.

Within the in vivo Mammalian Erythrocyte Micronucleus Test (OECD Guideline 474), the variables considered were:

-SPECIES (House mouse (as animal), Rat (as animal), Hamster (as animal), Human (as organism));

-STRAIN (Chinese or not reported in the case of Hamster; CD-1, ICR, Not reported, NMRI, Swiss, Tif:MAGf(SPF), Balb/c, B6C3F1, CF-1, C57BL, BDF1 mouse, SwissWebster, CBA, ddy mouse and CFLP in the case of House mouse; Fischer344, Wistar, Sprague-Dawley and Crj:CD(SD)in the case of Rat);

-NUMBER_INDIVIDUALS (from 3 to 130);

-SEX (Male/Female, Male and Female);

-ROUTE (oral: gavage, intraperitoneal, oral: unspecified, inhalation: unspecified, oral: feed, dermal, oral: capsule, intravenous and other);

-EXP_PERIOD (from 0.25 hour to 13 weeks);

-RESULTS (negative, positive, ambiguous and inconclusive);

-INVIVOTISSUEEXP (direct evidence-cytotoxicity, indirect evidence-systemic toxicity, direct evidence-cytotoxicity, no evidence, indirect evidence-systemic toxicity, indirect-toxicokinetic investigations and indirect evidence-systemic toxicity, indirect-toxicokinetic investigations).

1. opinion_pub_year (2005-2016)
2. literature_reference (332 documents)

We applied the RF technique to these 31 compounds with multiple in vivo micronucleus results to determine which of the above test variables might be more effective in discriminating between conflicting results. In other words, we used the RF technique to identify which of the test variables might influence the different results for each compound.

To perform the RF technique on the selected compounds, we used an *ad hoc* workflow developed using the KNIME platform (https://www.knime.com/). The core of the workflow is the Tree Ensemble Features node, which, along with the others, contains the RF Learner node.

### 6.2 Results

For 25 of these 31 compounds, the RF technique identified the most relevant variables associated with the different outcomes. SPECIES (for Triazole alanine); STRAIN (for 1-[(6-chloro-3-pyridinyl)methyl]-,oxohydrazone, 8-Hydroxyquinoline, Carbofuran, Carbosulfan, Dimethoate, Diuron, Glyphosate formulation (Roundup), Isoxaben and Sulcotrione); NUMBER_INDIVIDUALS (for 8-Hydroxyquinoline, Captan, Carbosulfan, Dazomet, Dimethoate and Diuron); SEX (for 1-[(6-chloro-3-pyridinyl)methyl]-,oxohydrazone, 2,4-D, Captan, Carbosulfan, Dazomet, Dimethoate, Isoproturon, Phosphine, Sulcotrione, Triazole alanine and Tribasic copper sulfate); ROUTE (for Carbendazim, Dazomet, Dimethoate and Propaquizafop); EXP_PERIOD (for 2,3,5,6-tetrachloroterephthalic acid); INVIVOTISSUEEXP (for 8-Hydroxyquinoline, Benfuracarb, Captan, Carbosulfan, Dazomet, Dimethoate, Ethofumesate, Phosphine and Propaquizafop); literature_reference (for 8-Hydroxyquinoline, Benfuracarb, Carbofuran, Carbosulfan, Cypermethrin, Eugenol, Glyphosate and Methyl isocyanate). Considering the number of compounds, the most relevant variables are SEX (11/25, 44% of coverage) INVIVOTISSUEEXP, STRAIN (9/25, 36% of coverage) and literature_reference (8/25, 32% of coverage).

In some cases, more than one variable may occur for the same compound. Indeed, the RF technique uses combinations of features in different trees. In some cases, a single variable is sufficient to discriminate against a class; in others, interactions between several features are required.

For the other six compounds (2-hydroxynaphthalene-1,4-dione, 5-(Trifluoromethyl)-2(1H)-pyridinone, 6-ethoxy-1,3,5-triazine-2,4-diamine, Tefluthrin, Tralkoxydim and Trifluralin) for which the RF could not explain the results, there were indeed no large variations in the values of the variables that could account for the differences.

Next, we extended the analysis and applied the RF technique to the other three subsets of data as previously mentioned, i.e. the Mammalian Bone Marrow Chromosome Aberration Test (OECD TG 475), the Mammalian Chromosome Aberration Test - OECD TG 473 and in vitro Mammalian Cell Gene Mutation Test - OECD TG 476. The detailed results are presented in supplement 3.

Among the four datasets, 122 compounds have multiple data. Nine of them (2,4-DB, 2,4-dichlorophenol, 2-(Misik et al.)-5-[(2RS)-2-(ethylsulfonyl)propyl]-3-hydroxycyclohex-2-en-1-one, 2-Phenylphenol, Bentazone, Bromuconazole, Cycloxydim, Dazomet, Prochloraz) are in common within the in vitro datasets meanwhile ten are in common within the in vivo datasets (Captan, Carbendazim, Carbofuran, Carbosulfan, Cypermethrin, Dimethoate, Glyphosate formulation (Roundup), Isoproturon, Methyl isocyanate, Propaquizafop). No compound is present in all four datasets.

Among the variables in common to the in vitro datasets, SPECIES and STRAIN have the highest value of coverage percentage (40% for the OECD 476). Among those in common to the in vivo datasets, NUMBER_INDIVIDUALS has the highest value of coverage percentage (50 % for the OECD 475). STRAIN, literature_reference and SPECIES are the variables in common to all the datasets, in vitro and in vivo.

### 6.3 Conclusion

We applied the RF technique to four different datasets of genotoxic endpoints to select the variables associated with the multiple outcomes and, therefore, most relevant for the uncertainty analysis. Among the in vivo methods, Mammalian Erythrocyte Micronucleus Test (*TG 474*) and Mammalian Bone Marrow Chromosome Aberration Test (*TG 475*) datasets were selected while among the in vitro methods, Mammalian Chromosome Aberration Test (*TG 473*) and Mammalian Cell Gene Mutation Test (*TG 476*) datasets were selected. For each dataset, those compounds showing multiple outcomes have been considered for the analysis. The utility of these results is discussed in section 9.

## 7. OECD TG conform study-variables that may drive study sensitivity

### 7.1 Introduction

To possibly develop recommendations for increasing the methods sensitivity, the results from the previous section were analysed in more detail.

It is clear from section 6 that several study variables may be important drivers of data variability for some compounds. To look closer into this, an additional analysis was done on the general difference in fraction of positives of compounds over a variable, such as strain, cell culture type (for in vitro assays), sex of the test subjects, route of exposure and the experimental period. This analysis assessed if the different options allowed in the OECD TGs for a particular variable are of significant influence for the test result, and the effect size.

### 7.2 Method

For this analysis, the same filtered dataset described in section 5 was used as the starting point (see also supplement 2a). The dataset was further filtered to include only those compounds that have at least one positive test in the assay of interest. Furthermore, ambiguous and inconclusive test results (neither positive nor negative) were excluded. If particular options within a variable were tested with relatively more genotoxic compounds, these would show a higher fraction of positive results. Therefore, an analysis was done that assessed if a particular option of a variable (e.g. strain) is of influence for the test result regardless of the fraction of genotoxic compounds tested. For this analysis, per assay, we divided the data into two groups, one with results of the option of the variable of interest (e.g. the option ‘CD-1’ in the variable ‘strain’) and one with results of all other options for the variable combined. Only compounds that are tested in both these groups were included in the analysis. This resulted in a selection of compounds with at least 2 replicates (not 3, as was done in section 5 and 6) distributed over two groups. The difference in fraction positives for these compounds was analysed between one group (consisting of the option of the variable of interest) and the group containing the rest of the options for the variable, with a paired Wilcox test. The Wilcoxon paired test is a nonparametric equivalent of the paired t-test. It calculates the difference between each pair of observations and ranks the absolute values of the differences. It then analyses the number of positive and negative differences and whether they are symmetric around zero. The Wilcoxon test is an alternative to the t-test when the normal distribution of the differences between paired individuals cannot be assumed.

### 7.3 Results

First, the variable ‘strain’ was analysed. For in vitro tests, cell culture was regarded as a similar variable as ‘strain’ in in vivo tests. To include both in vitro and in vivo assays, strain and cell culture are referred to as ‘test system’ in this analysis.

Table 7 shows, per OECD TG, to which extent the choice of the test system (strain or in vitro cell culture) results in higher sensitivity, this means a higher fraction of positives. In the database, many different options in test systems are available; however, only test systems with enough chemicals tested can be analysed.

**Table 7.** Significance of options in test systems (animal strains or in vitro cell culture) on the sensitivity (positive fraction) testing over compounds. Only results with enough tested chemicals to produce a p-value are shown.

| OECD TG | Test system (strain or in vitro cell culture) | Total substances | Mean difference of frequency positive tests between the test system and all other reported test systems | p-value |
| --- | --- | --- | --- | --- |
| 474 | CD-1 | 12 | -0.17 | 0.37 |
| 474 | NMRI | 3 | -0.06 | 0.75 |
| 474 | SWISS | 6 | 0.55 | 0.06* |
| 475 | SWISS | 6 | 0.93 | 0.03* |
|  | Sprague-Dawley | 4 | -0.58 | 0.13 |
| 473 | lymphocytes | 22 | -0.18 | 0.30 |
| 473 | CHO | 14 | -0.18 | 0.34 |
| 476 | lymphoma L5178Y cells | 19 | 0.25 | 0.18 |
| 476 | lung fibroblasts (V79) | 8 | -0.38 | 0.20 |

\* Significant $p < 0.1$ value

Table 7 shows that the test system ‘SWISS’ has a significant higher frequency of positive test results compared to other test systems as a group, for the same tested compounds.

A similar analysis was performed for the variable ‘sex’, but this was only possible for the two in vivo assays. The sex is reported in both assays as ‘male’, ‘female’, ‘male/female’, and ‘unknown’. Only very few, twenty-four and two of the reported tests, were labelled ‘female’ in assays 474 and 475. Therefore, the analysis was done on results reported as ‘male’ and ‘male/female’. This is suboptimal, however, if a significant difference is found, this indicates that testing only in ‘male’ may yield a different fraction of positive results (Table 8).

**Table 8.**
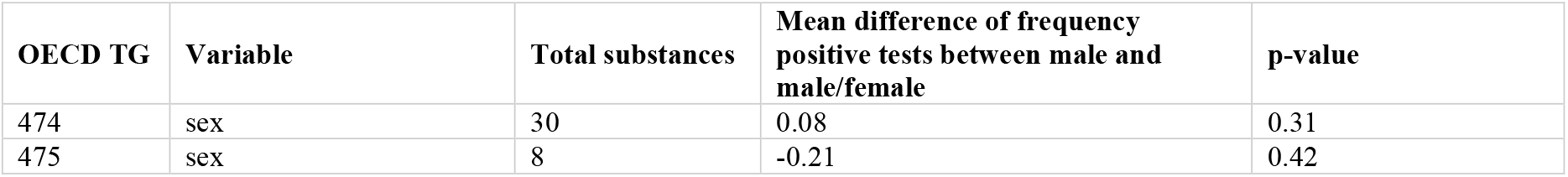
Significance of the variable ‘sex’ in testing per compound positive fraction. Only results with enough tested chemicals to produce a p-value are shown.

Table 8 shows that the difference was not significant in both in vivo assays. Moreover, the mean difference in frequency is negative in one assay and positive in the other. This implies that any influence of male vs. male/female testing on the frequency of positive tests is not systematic over assays.

The variable ‘Route’ was found as an important variable driving the test result in the RF analysis of section 6. This variable is also only relevant for the two in vivo assays. For the oral route, several variations are noted in the database: capsule, feed, gavage, unspecified. These were analysed as a single group.

Two routes dominate the database, intraperitoneal and oral. For this reason, the mean difference of frequency positive test results in Table 9 are nearly the same, but opposite (positive/negative). However, the results are not significant. Theoretically, more data could prove whether the route is a factor that drives sensitivity.

**Table 9.**
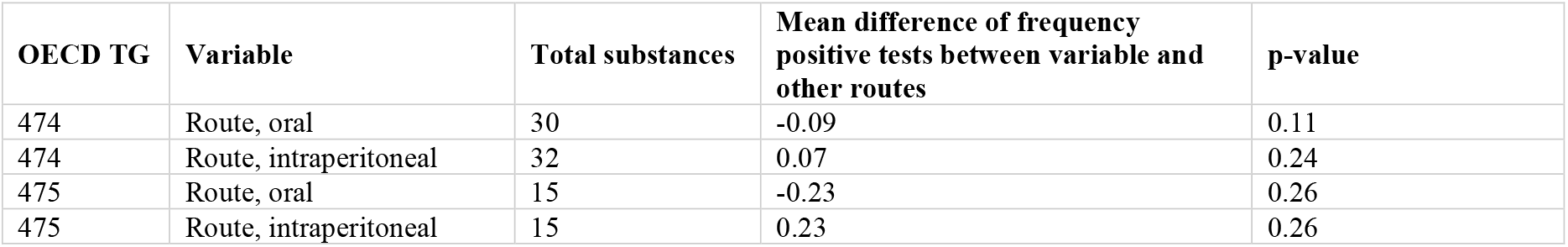
Significance of the variable ‘route’ in testing per compound positive fraction. Only results with enough tested chemicals to produce a p-value are shown. Some routes have few tests reported.

| OECD TG | Variable | Total substances | Mean difference of frequency positive tests between variable and other routes | p-value |
| --- | --- | --- | --- | --- |
| 474 | Route, oral | 30 | -0.09 | 0.11 |
| 474 | Route, intraperitoneal | 32 | 0.07 | 0.24 |
| 475 | Route, oral | 15 | -0.23 | 0.26 |
| 475 | Route, intraperitoneal | 15 | 0.23 | 0.26 |

Species is another variable between tests in the different assays.

Compounds tested in mouse tend to have slightly (10%) higher frequency of positives in *TG 474* compared to all other species. Although the difference is not significant in all TGs, this trend is repeated in the other TGs (Table 10).

**Table 10.** Significance of the variable ‘species’ in testing per compound positive fraction. Only results with enough tested chemicals to produce a p-value are shown.

| OECD TG | Variable | Total substances | Mean difference of frequency positive tests between variable and other species | p-value |
| --- | --- | --- | --- | --- |
| 474 | mouse | 19 | 0.1 | 0.02* |
| 475 | mouse | 16 | 0.16 | 0.21 |
| 473 | mouse | 4 | 0.63 | 0.17 |
| 473 | human | 39 | -0.11 | 0.26 |
|  | hamster | 41 | 0.05 | 0.53 |
| 476 | mouse | 52 | 0.09 | 0.18 |
|  | hamster | 52 | -0.09 | 0.20 |

\* Significant $p < 0.1$ value

For the different tests, experimental period was reported in hours, days, or weeks. This was recalculated to hours. The tests were divided into two groups (per assay): less or equal to the median experimental period (“short”) or higher than median experimental period (“long”).

Given that the mean difference in frequency of positive tests is very small between short and long experimental period, and the high p-values, this is not likely an important factor of influence, even if more data become available (Table 11).

**Table 11.** Significance of the variable ‘experimental period’ in testing per compound positive fraction. Only results with enough tested chemicals to produce a p-value are shown.

| OECD TG | Variable | Total substances | Mean difference of frequency positive tests between long and short experimental period | p-value |
| --- | --- | --- | --- | --- |
| 474 | Experimental period | 281 | 0.00 | 0.67 |
| 475 | Experimental period | 67 | 0.01 | 0.48 |

The addition of metabolic enzymes (‘MET_INDICATOR’) is a variable relevant only to in vitro assays. Table 12 shows the test results for the difference in sensitivity between tests with and without the addition of these enzymes.

**Table 12.**
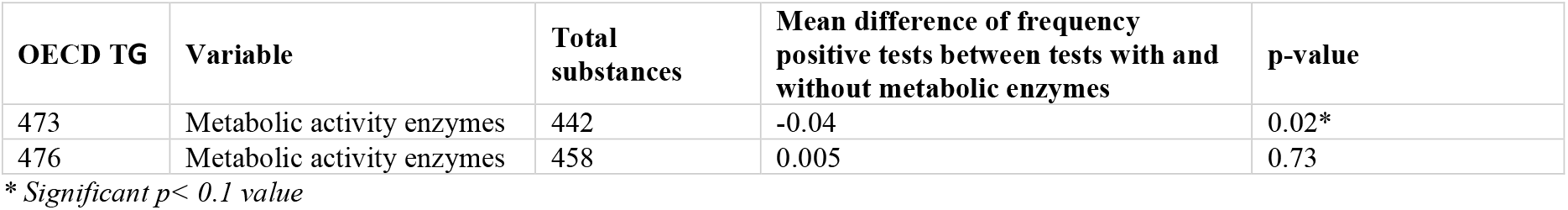
Significance of the variable ‘metabolic activity enzymes’ in testing per compound positive fraction. Only results with enough tested chemicals to produce a p-value are shown.

| OECD TG | Variable | Total substances | Mean difference of frequency positive tests between tests with and without metabolic enzymes | p-value |
| --- | --- | --- | --- | --- |
| 473 | Metabolic activity enzymes | 442 | -0.04 | 0.02* |
| 476 | Metabolic activity enzymes | 458 | 0.005 | 0.73 |

\* Significant $p < 0.1$ value

Although the p-value is significant for a difference in sensitivity (which seems lower with the addition of enzymes), the effect size is small, and the effect is not consistent between the two in vitro assays. Therefore, the addition of metabolic enzymes is not a likely structural influence for inducing different sensitivity in genotoxicity testing. Note, that mechanistically it is clear that metabolic activation may significantly increase the sensitivity of the tests for specific compounds. Yet, among the 442 or 458 compounds (for *TG 473 or 476*) with at least two replicates and at least one of them positive, this was statistically not apparent. One may speculate that more replicate results are available for compounds with less clearly positive results.

The last variable that showed significant influence on genotoxicity testing in the RF analysis was the number of individuals. It can be hypothesized that a test is labelled ‘positive’ more frequently if more individuals are tested. The tests were divided into two groups (per assay): less than median number of individuals (“low”) or higher than median number of individuals (“high”).

The analysis shows that the number of individuals used in the test is not significantly associated with the frequency of genotoxicity. Moreover, the mean difference in frequency is negative in one assay and positive in the other (Table 13).

**Table 13.** Significance of the variable ‘number of individuals’ in testing per compound positive fraction. Only results with enough tested chemicals to produce a p-value are shown.

| OECD TG | Variable | Total substances | Mean difference of frequency positive tests between high and low number of individuals | p-value |
| --- | --- | --- | --- | --- |
| 474 | Number of individuals | 17 | 0.12 | 0.1 |
| 475 | Number of individuals | 6 | -0.25 | 0.37 |

**Table 14.** Variability information published from other fields of (eco)toxicology.

|  | Methods | Similarity of (pseudo-) replicate results | References, reviewed & summarized in (Paparella et al. 2020; Paparella et al. 2017; Paparella et al. 2013); |
| --- | --- | --- | --- |
| Categorical results | AMES test | ca. 90% (on average for chemicals) for 2 categories | (Benigni and Giuliani 1988) (42 compounds, 12 labs); (Piegorsch and Zeiger 1991) (209 compounds); (Sushko et al. 2010) (1680 compounds) |
|  | LLNA | 70-80% for 2 categories<br>60-70% for 3 categories | (Dumont et al. 2016); (Hoffmann et al. 2018); (Hoffmann 2015); (Roberts et al. 2016); |
|  | Eye damage |  | (Adriaens et al. 2014); (Barroso et al. 2017) |
|  | 2-year rodent carcinogenicity | 57% for 2 categories<br>25-40% discordant between rats and mice | (Gottmann et al. 2001) (121 chemicals, NCI/NTP database vs. literature papers); (Alden et al. 2011), (Brambilla et al. 2012) (Friedrich and Olejniczak 2011), (Van Oosterhout et al. 1997) (110 to 181 drugs) |
|  | Repeated dose toxicity studies | Qualitatively discordant > 20% (e.g. increasing vs. decreasing, no-response vs. response, even when known rat-specific effects were not taken into consideration); from 880 available data sets: 58 substances, 91 pairs of studies, 2 sexes, 6 endpoints (body weight, liver & kidney weight, erythrocytes) | (Bokkers and Slob 2007) |
|  | Repeated dose toxicity studies (ToxRefDB) | Qualitative reproducibility of organ-level effects in repeat dose studies of adult animals was 33-88%, depending on grouping. (Organs associated with more negative chemicals (stomach, thyroid, adrenal) had higher rates of concordance. Within species concordance tended to be greater than within-study concordance.) | (Friedman et al. 2023) |
| Continuous results | Repeated dose toxicity studies (ToxRefDB) | Replicate LOAELs may vary by factor 100 | (Pham et al. 2020) |
|  | Acute fish toxicity LC50 | Replicate LC50s may range several orders of magnitude, with CVs typically between 50% and 100%, up to > 400% | (Schur et al. 2025) (805 compounds) |
|  | Motor-activity in rats | LOAEL ratio range = 1-6 | (Crofton et al. 1991) (9 chemicals in 6 labs) |
| | Behavioural tests | LOAELs varied by factor $\geq 10$ | (Hunter et al. 1979) |

This suggests that any influence of the number of individuals on the frequency of positive tests is not systematic across assays. It may be that negative results occasionally lead to the inclusion of more animals within a confirmatory test to ensure reliability while a positive result is interpreted as definitive. This could create an association where tests with larger sample sizes exhibit a lower fraction of positive results. However, it is unknown if this is the case for the analysed data set.

### 7.4 Conclusion

The analyses indicate that variables strain and/or species could, in theory, be drivers of sensitivity. The ‘Swiss’ strain yielded a higher frequency of positive tests compared to other strains. This is an often-used mouse strain, and species ‘Mouse’ concurrently yielded a higher frequency of positive tests than other species. The species effect (mouse) was significant in some of the four analysed assays and the effect size was univocal. Because most used mice are of Swiss strain, these variables cannot be seen separate. Route (oral or intraperitoneal) in in vivo assays could be a driver as well; it was near significant and effect size was univocal. Though intraperitoneal yielded a slightly higher frequency of positive tests, this may not be a useful statistical result, since it is today considered as an unrealistic exposure route. The future availability of more test results could increase statistical power to settle this with more certainty. Not likely or unequivocal drivers of sensitivity are number of individuals used, sex, addition of metabolic enzymes and experimental period. Sex, and number of individuals used, had opposite effect size between both in vivo assays. Addition of metabolic enzymes had (small) opposite effect sizes between both in vitro assays. Experimental period was nowhere near significant with a near-zero effect size. The interaction of effects was not tested. In theory, however, it could be that variables are confounding and that correcting for mixed (random and fixed) effects can give a more accurate result. Additional data, from a large-scale experiment with a crossed design could distinguish and correct for the influence of the different variables that determine the frequency of positive tests for chemicals. If feasible at all, such work towards methods improvement would be much easier for in vitro methods, considering costs, practicalities and in the current European efforts for a roadmap towards an animal-free regulatory system. The meaning of these results for the purpose of this paper is further discussed in section 9.

## 8. How does the variability summarized here for Genotoxicity TG data compare to the variability reported in other fields of (Eco)toxicology?

Similarity of pseudo-replicate results was analysed for methods from other fields of (eco)toxicology. Such similarity ranged from 33% to 90% for categorical results, and more than one order of magnitude for continuous results, depending on the method type. Within this manuscript on genotoxicity, we provide a variability analysis with a higher resolution, i.e. in terms of percent compounds within pseudo-replicate similarity ranges of < 66%, 66-85% and > 85%. Within this analysis, the (pseudo-)replicate similarity appeared to be below 85% for 22% to 77% of the compounds, depending on the genotoxicity TG (see section 5). Due to the different resolution, these variability values are not directly comparable with those presented in table 13. Yet, it appears obvious that the variability of results from standardized methods observed in this analysis is not a specific challenge for genotoxicity methods, but important and critical for many methods.

It is noted that estimates for the similarity of (pseudo-)replicate categorical results may also be biased by the number of compounds close to the category boundaries. Within databases containing a higher number of compounds close to the category boundaries, the variability in the methods readouts will more strongly impact categorical results variability. A more sophisticated approach, excluding a borderline range between the categories was applied for skin sensitization (Leontaridou et al. 2017). Such an improved variability assessment is beyond the resources available for the analysis presented here. However, it is noted that ultimately, when potency read-outs from genotoxicity methods will become available and accepted, the respective data variability assessment shall be improved.

## 9. Conclusion: How this knowledge on variability and uncertainties may be used for the NAM based IATA development

The NAM-based IATA may be developed considering sensitivity and specificity estimates for the individual NAMs and for some combinations thereof. Such accuracy estimates may be based on genotoxicity calls from chemical reference data sets. However, genotoxicity TGs have been improved over time, such that databases contain data with variable quality (see section 2) and genotoxicity calls may greatly differ between databases, probably due to inclusion of datasets which differ in data number per compound and data quality, and clearly due to different data interpretation criteria applied within the different databases (see section 3). Therefore, attributing genotoxicity calls to reference chemicals is a complex task and necessarily uncertain. Ideally, an Expert Knowledge Elicitation (EKE) approach could be used (EFSA 2014): In this case, a group of experts would individually review the heterogeneous datasets and provide their expert specific calls for each of the compounds. Using a moderated discussion, the arguments for these calls could then be exchanged between the experts. Finally, after discussion, the uncertainty of the calls and the resulting uncertainty of the accuracy estimates for the methods could be expressed as a democratically integrated probability distribution of expert opinions. Yet, such a process was not yet applied for generating genotoxicity calls.

Moreover, such an EKE would not discern the variability of expert judgments (stemming from a variable weighting of the different data) from biological data variability. An estimate for the latter is provided in section 5 based on the EFSA genotoxicity database. It appears that more than 50% of the compounds tested have a chance of less than 86% to show similar genotoxicity calls upon replication within the in vivo micronucleus test, the in vivo bone marrow chromosomal aberration test but also within the in vitro mutation and in vitro chromosomal aberration test. For the AMES test, this low chance for similar results appears true for fewer, i.e. about 20%, of the compounds. Due to the stringent study quality filters applied, i.e. “acceptable study, according, equivalent or similar to the TG”, too few data remain for a variability estimate for the other TGs.

Genotoxicity results have so far mainly been used for hazard identification rather than hazard characterisation. Nevertheless some compounds may have a potency that is borderline to be detectable with the current tests, such that the number of such compounds in the dataset may influence variability estimates for the methods. However, the potency is not reported within the EFSA genotoxicity database, currently, this uncertainty cannot be assessed. Moreover, the variability estimate is uncertain for data from current TG versions, since it includes also older data, which were acceptable for regulation at the date of the EFSA opinion (2004-2016). Since science and methods will continue to evolve, data heterogeneity will likely continue to remain a challenge for any statistical analysis requiring ample datasets.

However, the EFSA publication date did not statistically explain the variability in results from individual compounds. Interestingly, literature reference explained data variability only for about 30% to 40% of the compounds. This indicates that variability is likely to be dominated by the allowed protocol differences in the TGs rather than by the reproducibility (of identical protocol variants) in the strict sense. Most important variables for statistically explaining variability appear to be species and strain for all analyzed methods, number of individuals, exposure route, confirmation of exposure and sex for in vivo methods and metabolic activation for in vitro methods. Theoretically, such information may be used to optimize method standardization (section 6). However, it would take considerable resources and time to validate such optimization and to improve the availability of data with low variability. Also, it would need further consideration if reducing variability may negatively impact the relevance of the results, since it may be difficult to define which is the most human-relevant test system in terms, of e.g. strain, sex or exposure route.

For statistically explaining differences in overall sensitivity for compounds in general, between options that are allowed in variables in OECD TGs, the variables species, strain and exposure route appear to be most important (section 7). More specifically, the species ‘mouse’, the strain ‘Swiss’ and exposure route ‘intraperitoneal’ were more sensitive than other options within the variables. However, new data from a large-scale experiment with a crossed design would be necessary to distinguish and correct for the influence of the different variables. If at all feasible, such work might be practically possible for in vitro methods only. While it is important from a precautionary perspective to increase the sensitivity of a method, the optimum of sensitivity may not necessarily be maximum sensitivity, but relevant sensitivity. Within current regulatory guidance for chemicals and biocides, in vitro methods are considered rather too sensitive and therefore may be overruled with negative in vivo data. In any case, at this moment, any attempt to stratify the available data for such a refined analysis of sensitivity would result in too few data for a robust sensitivity analysis.

As a consequence, the uncertainties about potential drivers of variability and sensitivity need to be considered qualitatively. In the moment, we can neither identify them with a high reliability, nor clearly define what their optimum would be. Moreover, there are significant practical limitations to improving respective knowledge. Thus, recognizing and accepting a certain degree of unavoidable uncertainty in biological data and their conclusions appears crucial for advancing integrated testing and assessment methods. Notably, such uncertainty is inherent in the current animal data-based approaches as well as non-animal-methods based approaches.

However, since the data variability limits possible data correlations between different methods, an estimate for data variability from similar methods may be used as a benchmark to define equivalent results. Based on the qualification and quantification of uncertainty provided here, data correlations between different methods beyond 85% are not likely to be real but rather due to a limited data availability or selection bias. Data correlations between 66% and 85% are more likely to be within the methods variability range. Such an estimate would also fit within the variability ranges identified for other toxicological methods (section 8).

Consequently, for the development of a NAM-based IATA any accuracy estimate for NAMs relative to reference data sets needs to be considered with great caution. Low variability and mechanistic relevance and complementarity may be more robust criteria for integrating an intelligent combination of NAMs.

## Acknowledgement

This work was carried out in the framework of the European Partnership for the Assessment of Risks from Chemicals (PARC) and has received funding from the European Union’s Horizon Europe research and innovation programme HORIZON-HLTH-2021-ENVHLTH-03-01 under Grant Agreement No 101057014. Views and opinions expressed are, however, those of the author(s) only and do not necessarily reflect those of the European Union or the Health and Digital Executive Agency. Neither the European Union nor the granting authority can be held responsible for them. This work was further co-funded by all institutions affiliated to the co-authors and in addition for Martin Paparella by the Austrian Federal Ministry for Environment, Department V/5 - Chemicals Policy and Biocides.

The authors would like to express their sincere gratitude to Juan Manuel Parra Morte from the European Food Safety Authority for his valuable guidance and expert advice in interpreting the EFSA genotoxicity database.

## Ethical standards

The manuscript does not contain clinical studies or patient data.

